# Inferring the correlated fitness effects of nonsynonymous mutations at the same site using triallelic population genomics

**DOI:** 10.1101/029546

**Authors:** Aaron P. Ragsdale, Alec J. Coffman, PingHsun Hsieh, Travis J. Struck, Ryan N. Gutenkunst

## Abstract

The distribution of mutation fitness effects is central to evolutionary genetics. Typical univariate distributions, however, cannot model the effects of multiple mutations at the same site, so we introduce a model in which mutations at the same site have correlated fitness effects. To infer the strength of that correlation, we developed a diffusion approximation to the triallelic frequency spectrum, which we applied to data from D. melanogaster. We found a moderate correlation between the fitness effects of nonsynonymous mutations at the same codon, suggesting that both mutation identity and location are important for determining fitness effects in proteins. We validated our approach by comparing with biochemical mutational scanning experiments, finding strong quantitative agreement, even between different organisms. We also found that the correlation of mutation fitness effects was not affected by protein solvent exposure or structural disorder. Together, our results suggest that the correlation of fitness effects at the same site is a previously overlooked yet fundamental property of protein evolution.

---

Mutation effects on fitness range from strongly deleterious to strongly beneficial, and the distribution of mutation fitness effects (DFE) is key to many problems in genetics, from the evolution of sex [1] to the architecture of human disease [2]. In general, there are many strongly deleterious mutations, a similar number of moderately deleterious or nearly-neutral mutations, and a small number of beneficial mutations [3]. The DFE may be determined experimentally through direct measurements of mutational fitness effects in clonal populations of viruses, bacteria, or yeast [4, 5], and recent studies have provided high resolution DFEs for single genes [6, 7] and for beneficial mutations [8]. The DFE may also be inferred from comparative [9, 10] or population genetic [11, 12, 13, 14] data, although this approach has little power for strongly deleterious mutations. In the typical population genetic approach, the population demography is first inferred using a putatively neutral class of mutations and the DFE for another class of mutations is inferred by modeling the distribution of allele frequencies expected under a model of demography plus selection.

Most population genetic inference has focused on biallelic loci, where the ancestral and one derived allele are segregating in the population [15]. When many individuals are sequenced, however, even single-nucleotide loci are often found to be multiallelic, with three or more segregating alleles. Multiallelic loci pose a challenge for modeling selection. To use a typical univariate DFE, one must assume that mutations at the same site either all have equal fitness effects (so that mutation location completely determines fitness) or independent fitness effects (so that mutation identity completely determines fitness). Neither of these assumptions is biologically well-founded, suggesting the need for more sophisticated models of fitness effects. Here we introduce a model of correlated fitness effects for mutations at the same site, and we analyze sequence data to infer the strength of that correlation.

Our inference is based on triallelic codons, sites where three mutually nonsynonymous amino acid alleles are segregating in the population (Fig. 1A). Hodgkinson and Eyre-Walker recently found in humans a roughly two-fold excess of triallelic sites over the expectation under neutral conditions and random distribution of mutations [16]. This led them to suggest an alternate mutational mechanism that could simultaneously generate two unique mutations, although recent population growth and substructure can account for the distribution of observed triallelic variation [17]. Recently, Jenkins, Mueller and Song developed a coalescent method to calculate the expected triallelic frequency spectrum under arbitrary single-population demography. They showed that triallelic frequencies are sensitive to demographic history [18, 17], but their method cannot model selection.

**Figure 1:**
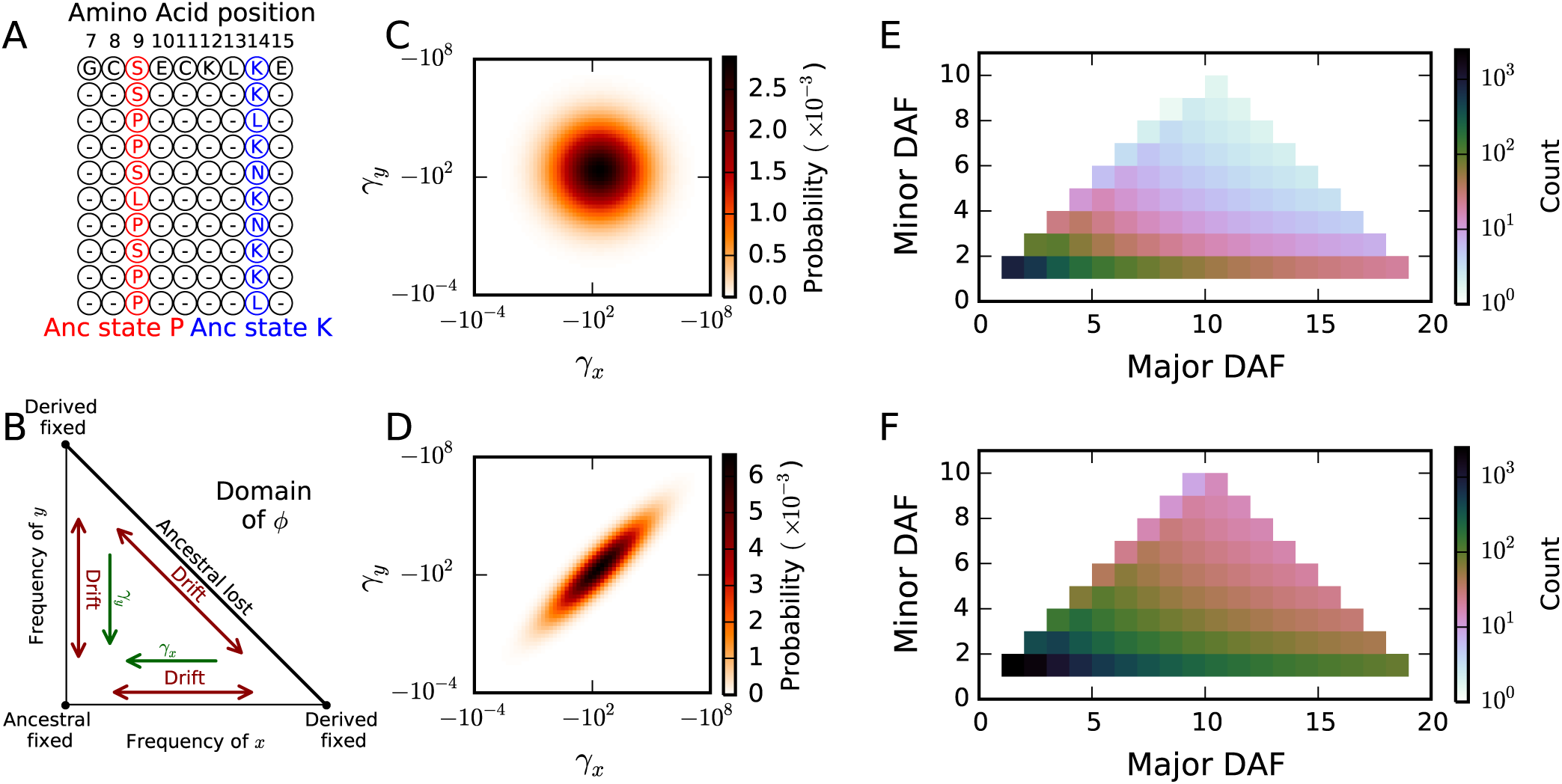
The triallelic frequency spectrum. A: Mutually nonsynonymous triallelic loci in protein coding regions have three observed segregating amino acid alleles. Here, with ten sampled chromosomes, at position9 the major and minor derived alleles, Serine (S) and Leucine (L), have frequencies 4 and 1, so this site contributes to the (4,1) bin of the TFS. Similarly, position 14 contributes to the (2,2) bin. B: The domain of the triallelic diffusion equation, *φ*, from Equation 1. The corners correspond to fixation of one of the three alleles, and the edges correspond to loss of one of the three alleles. New mutations enter the population along the horizontal and vertical axes, with density dependent on the background biallelic frequency spectrum. Pairs of selection coefficients for the two derived nonsynonymous mutations are sampled from a bivariate DFE, which includes a parameter for correlation between selection coefficients *ρ*. C: For an uncorrelated DFE, with *ρ* = 0, the selection coefficients are independent and often dissimilar. D: For strong correlation, here *ρ* = 0.9, selection coefficients are typically very similar. E, F: The correlation coefficient affects the expected frequency spectrum, with stronger correlation (F: *ρ* = 0.9) resulting in a higher proportion of intermediate- to high-frequency derived alleles and more triallelic sites overall relative to weak correlation (E: *ρ* = 0).

We developed a numerical diffusion simulation of expected triallelic allele frequencies including selection, and we coupled that simulation to a DFE that models the correlation between fitness effects of the two derived alleles. We applied this approach to infer the correlation coefficient of fitness effects from whole-genome *Drosophila melanogaster* data [19], inferring a moderate correlation between fitness effects of nonsynonymous mutations at the same site. To validate our inference, we compared with direct biochemical experiments, finding strong agreement. Lastly, we applied our approach to biologically relevant subsets of nonsynonymous mutations, to assess how the fitness effects correlation varies among classes of mutations.

## Results and Discussion

### The triallelic frequency spectrum with selection

The triallelic frequency spectrum summarizes sequence data from a sample of individuals by storing the counts of triallelic loci with each set of observed derived allele frequencies [17] (Fig. 1E,F). Because the order in which the two derived alleles arose is often unknown, we used counts of major and minor derived alleles, which have respectively higher or lower sample frequencies.

To obtain the expected sample frequency spectrum for a given model of selection and demography, we numerically solved the corresponding diffusion equation. First described by Kimura [20, 21, 22], the triallelic diffusion equation models the evolution of the density function *φ*(*x, y*) for the expected number loci in the population with derived allele frequencies (*x*, *y*), with *x*, *y* ∈ (0,1) and *x* + *y* < 1 (Fig. 1B):

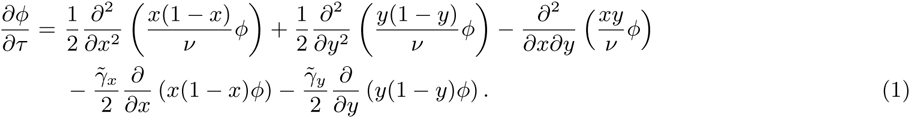

Time *τ* is measured is units of 2*N_a_* generations, where *N_a_* is the ancestral effective population size. The spatial second derivative terms account for genetic drift, which is scaled by the relative population size *ν*(*τ*) = *N*(*τ*)/*N_a_*, and the mixed derivative term accounts for the covariance in allele frequency changes. The population-scaled selection coefficient is *γ* = 2*N_a_s,* where s is the relative fitness of the derived versus ancestral allele. Here that selection coefficient must be adjusted 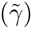 to account for interaction between selection acting on the two derived alleles (see Methods). Triallelic loci are created when a novel mutation occurs at a site that is already biallelic. The new allele initially has frequency 1/2*N*, and the existing derived allele has a frequency *x* (or *y*) ∈ (0,1) drawn from the population distribution of biallelic allele frequencies.

Some analytic results are known for triallelic diffusion [23, 24, 25], but we solved Eq. 1 numerically. We used a finite-difference method similar to that in ∂a∂i [26], although the mixed derivative term and the *x* + *y* < 1 boundary introduce complications (Fig. S1, Methods). We directly coupled with ∂a∂i to track the distribution of biallelic derived allele frequencies needed for creating triallelic loci. To obtain the expected sample frequency spectrum, we integrated over the population spectrum *φ* using trinomial sampling (Methods). We validated our numerical solution by comparison to the neutral coalescent solution (Fig. S2) and Wright-Fisher simulations with selection (Fig. S3).

Because there are two derived alleles, the DFE for triallelic sites is a two-dimensional joint distribution. We used a bivariate lognormal model for the DFE (Fig 1C,D). Because we assumed no knowledge of which allele arose first, the two marginal distributions are identical. The correlation between selection coefficients is then characterized by the correlation coefficient *ρ*. If *ρ* = 0, the selection coefficients of the two derived alleles at a single triallelic locus are independent, whereas if *ρ* = 1, they are equal. For a fixed marginal DFE, as the correlation coefficient *ρ* increases, more segregating triallelic loci are expected, particularly at moderate and high derived-allele frequencies (Fig. 1C-F). We quantified the relative importance of identity and location for protein mutation fitness effects through *ρ*; low correlation suggests that identity is more important, whereas high correlation suggests that location within the protein is more important.

### Correlation of selection coefficients for nonsynonymous mutations at the same site

To estimate the correlation between fitness effects of amino-acid altering mutations, we used 197 Zambian *D. melanogaster* whole genome sequences from Phase 3 of the *Drosophila* Population Genomics Project (DPGP3) [19]. We chose this population because it has high genetic diversity (and thus many triallelic sites) and a relatively simple demographic history [19]. We first modeled demographic history using biallelic synonymous sites. We then inferred the marginal DFE for newly arising nonsynonymous mutations using that demographic model and the biallelic nonsynonymous data. Lastly, we inferred the fitness effects correlation coefficient using our inferred demography and marginal DFE and the mutually nonsynonymous triallelic loci in the data.

We used ∂a∂i [26] to fit a three-epoch population size model to the unfolded biallelic synonymous frequency spectrum (Fig. 2A-B, Table S1). For all model fits, we included a parameter to account for ancestral state misidentification, which creates an excess of high-frequency derived alleles [27]. We fixed this demographic model for all future inferences, and we fit a univariate DFE to the biallelic nonsynonymous data. For negatively-selected sites (*γ* < 0), we assumed a lognormal distribution of selection coefficients with mean and variance parameters *μ* and *σ*, which has been previously shown to adequately describe the biallelic DFE for *D. melanogaster* [28]. Our DFE also included a point-mass modeling a proportion *p*_+_ of positively selected sites with scaled selection coefficient *γ*_+_. Our inferred biallelic DFE (Fig. 2C, Table S1) fits the data well (Fig. 2A), with just under 1% of new mutations inferred to be beneficial (inferred *γ*_+_ = 39.9). When fitting the DFE to the nonsynonymous data, the parameters for the lognormal portion (negatively selected sites) were tightly constrained, but *p*_+_ and *γ*_+_ were confounded and inversely correlated, as found in other studies [29, 30]. Our inferred proportions of mutations in various selective regimes agreed well with prior work (Table S2).

**Figure 2:**
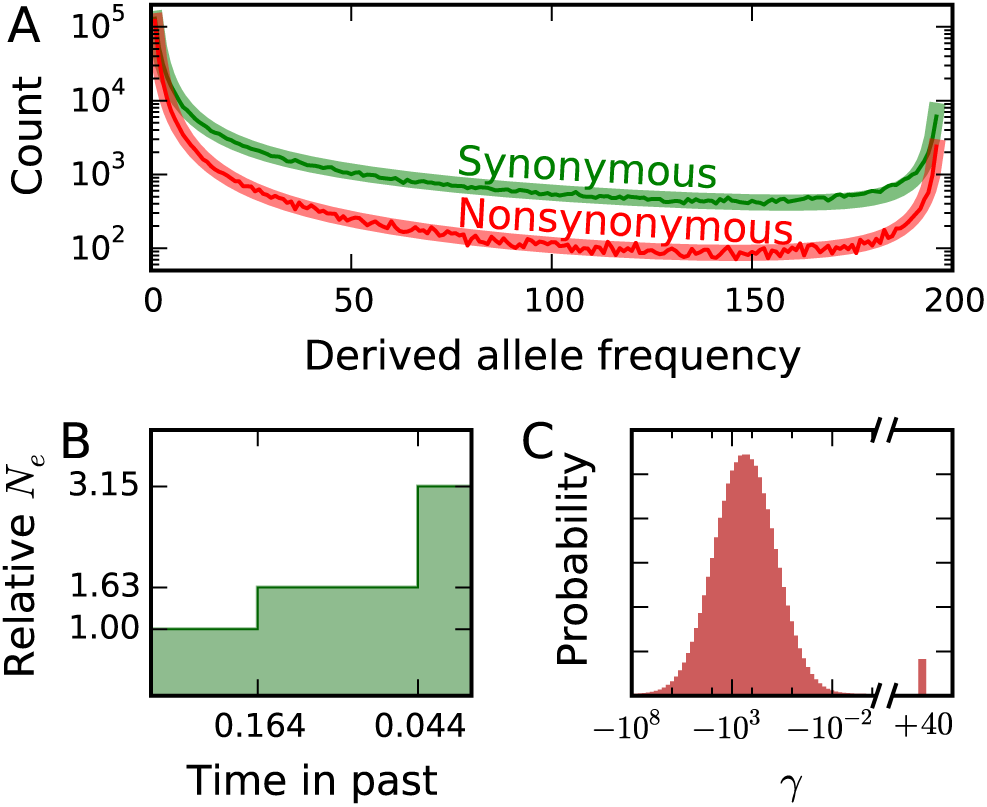
Inferences of demographic history and marginal distribution of fitness effects from biallelic data. A: Biallelic synonymous and nonsynonymous data (solid lines) and corresponding maximum-likelihood model fits (shaded). Both models include a parameter to account for ancestral state misidentification, which is likely responsible for most of the excess of high-frequency derived alleles. B: Inferred demographic model, with two instantaneous population size changes. C: Inferred distribution of fitness effects, lognormally distributed for negatively selected mutations with a proportion of positively selected mutations.

We worked at the codon level to assess the correlation in selection coefficients for nonsynonymous mutations, so a triallelic locus could arise from two mutations at the same nucleotide or at different nucleotides in the same codon. We extended our inferred one-dimensional DFE to two dimensions, fixing the parameters *μ*, *σ, γ*_+_, and *p*_+_, so that the correlation coefficient *ρ* was the only free parameter of the bivariate lognormal distribution, along with a single parameter for ancestral misidentification. Fitting to 10,471 mutually nonsynonymous triallelic loci (Fig. 3A), we inferred *ρ* = 0.51 (Fig. 3B, Table 1). Selection coefficients for nonsynonymous mutations at the same codon are thus somewhat but not completely correlated, so location and identity play roughly equal roles in determining mutation fitness effects.

**Figure 3:**
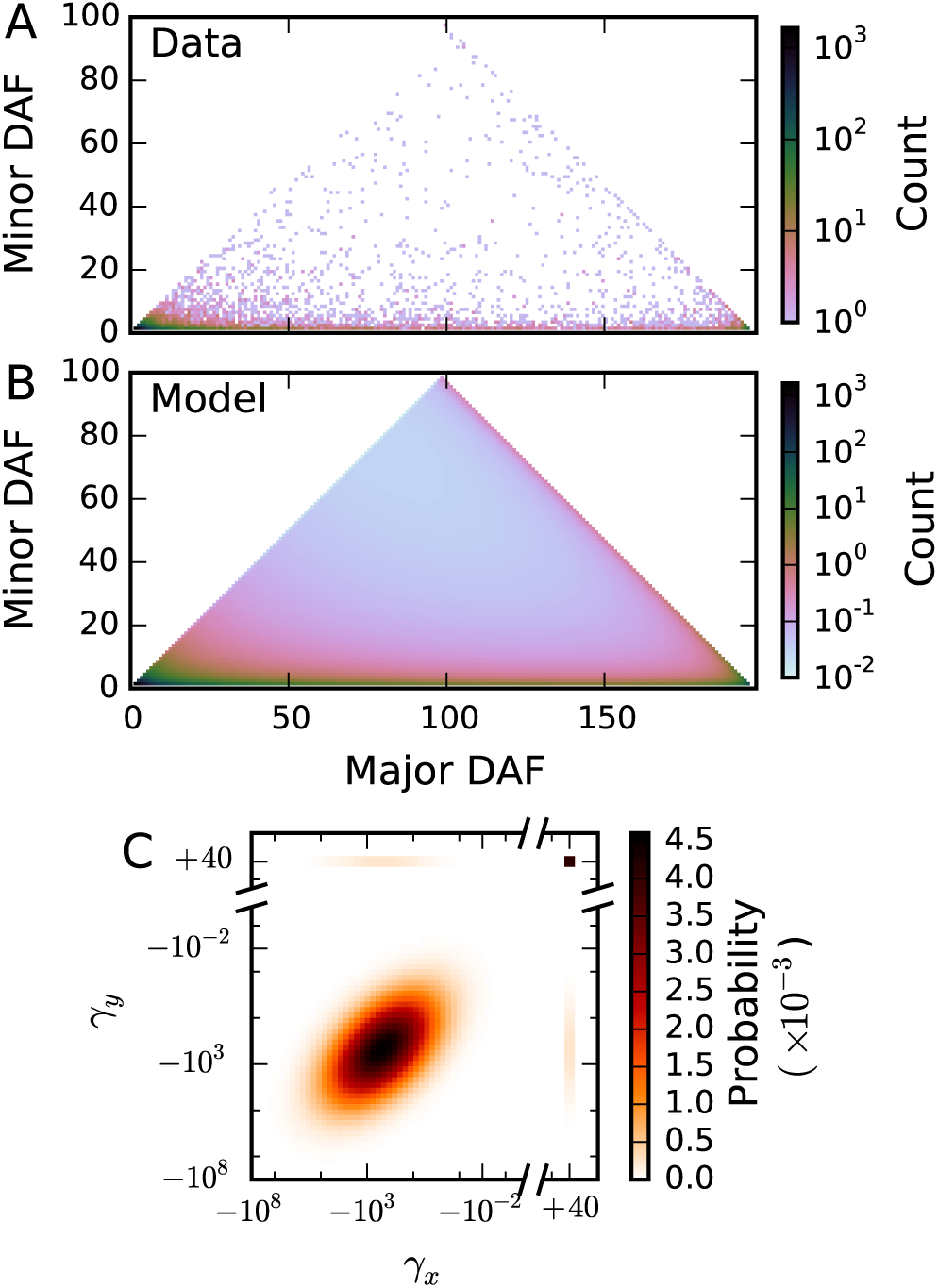
Inference of selection correlation coefficient from triallelic data. A: The observed triallelic frequency spectrum for mutually nonsynonymous triallelic sites, which contained 10,471 triallelic sites. B: The best fit model, optimizing the correlation coefficient *ñ* and the ancestral misidentification parameters. C: Joint distribution of selection coefficients from the maximum likelihood inferred correlation coefficient of *ñ* = 0:51. Selection coefficients for nonsynonymous mutations at the same site are moderately correlated.

**Table 1:**
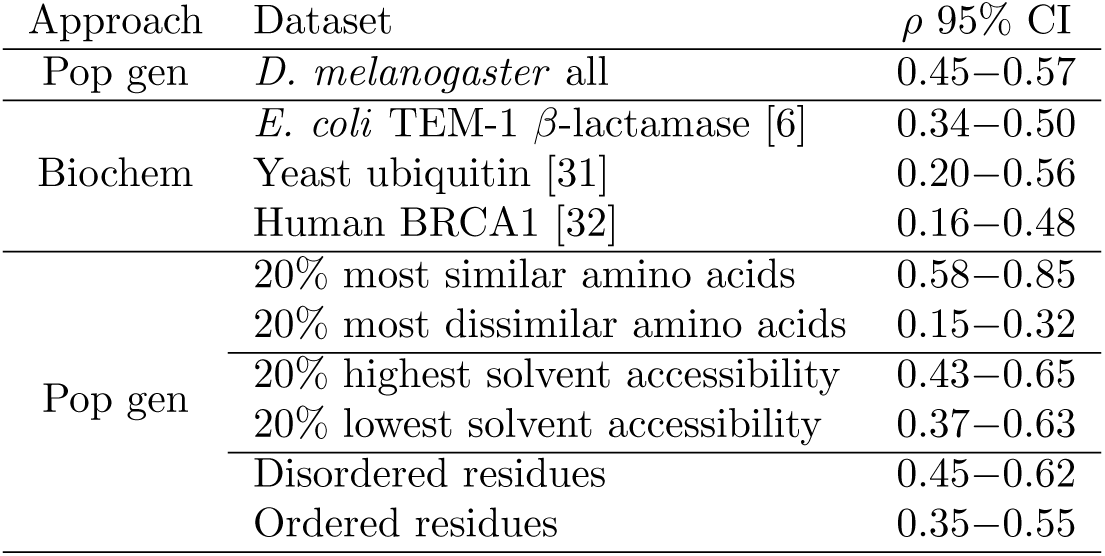
Fitness effect correlation coefficients for nonsynonymous mutations at the same codon, inferred from population genomic data and biochemical experiments.

### Comparison to experimental mutation scanning studies

Our population genetic approach allows us to simultaneously study the whole genome, but it is an indirect approach to measuring the selection coefficient correlation. Complementary experimental data come from mutational scanning experiments, which use deep sequencing to simultaneously assay the function of thousands of mutant forms of a protein [33] (Fig. 4A). To measure selection coefficient correlations from such data, we sampled pairs of mutually nonsynonymous mutations for each site assayed in the protein and calculated the resulting correlations (Fig. 4B, Supporting Text). Because our population genetic inference is insensitive to strongly deleterious mutations, we restricted our analysis to the moderately deleterious mutations found in each experiment (Fig. S4). We analyzed proteins from *E. coli* [6], *S. cerevisiae* [31], and humans [32] (Table 1). In all three cases these direct biochemical assays yielded a fitness effects correlation in agreement with our population genetic estimate, although the limited number of sites within each experiment yielded large confidence intervals, and experimental noise would tend to systematically bias the experimental correlations downward. These results suggest that the moderate correlation of mutation fitness effects we found in *D. melanogaster* also holds true for other organisms and proteins.

**Figure 4:**
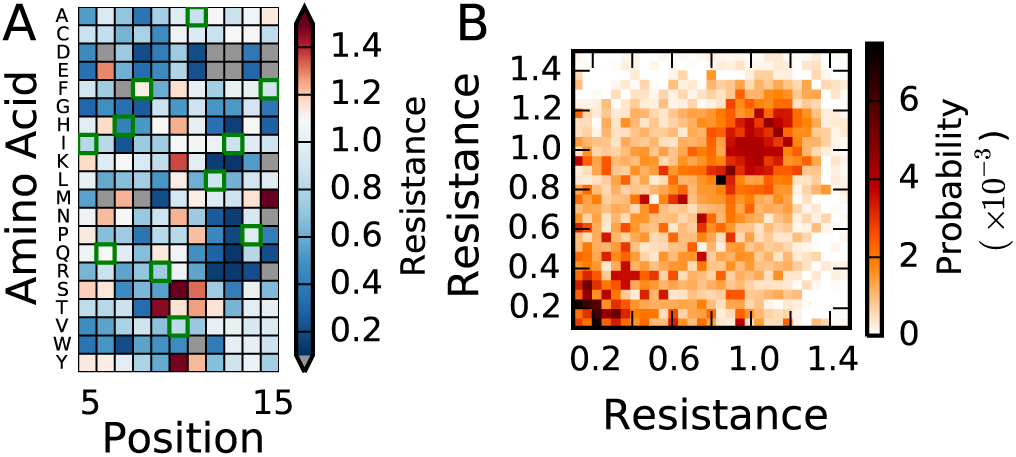
Mutational scanning data. A: Partial mutational fitness landscape for *E. coli* TEM-1 *â*-lactamase, adapted from [6]. For almost all possible single mutants, fitness was assayed as relative antibiotic resistance. Gray entries denote mutations not measured, and green squares highlight the ancestral sequence. B: Joint distribution of fitnesses for mutually nonsynonymous mutations for TEM-1 *â*-lactamase, using data from [6]. For other data sets in Table 1, see Fig. S4.

### Selection coefficient correlation for subsets of data

Sites within proteins vary in their evolutionary properties [34, 35], so we asked how the fitness effect correlation coefficient differs among subsets of the *D. melanogaster* population genomic data. We first tested our expectation that biochemically similar derived amino acids would have more tightly correlated selection coefficients than dissimilar derived amino acids [36, 37]. We assessed similarity using the Grantham matrix [38], which scores pairs of amino acids based on similarity of biochemical properties. We then refit the correlation coefficient and misidentification parameter to the subsets of loci with the top and bottom 20% of similarity scores. We indeed found that highly similar derived amino acids exhibited stronger correlation than dissimilar amino acids (Table 1), validating our approach.

We also assessed the correlation of fitness effects for subsets of amino acids that are buried or exposed, based on solvent accessibility, as well as subsets that are ordered or disordered, because protein structural properties are known to affect the amino acid substitution process [39]. We used SPINE-D [40] to separate sites into the top and bottom 20% of solvent accessibility scores and into disorder and ordered classes. For each subset, we refit the underlying marginal DFE and then fit the bivariate DFE to measure the correlation coefficient. As expected [41, 42, 43, 44], for buried residues with low solvent accessibility and for ordered residues, we inferred DFEs that were more negatively skewed than for residues with high solvent accessibility or that were structurally disordered (Table S3). We found, however, that these structural features did not affect the inferred fitness effects correlation coefficient (Table 1). Together, these results suggest that models of protein evolution that incorporate structural features [45, 46], do need to account for differences in the marginal DFE, but not for differences in correlation.

## Conclusions

We developed a novel numerical solution to the triallelic diffusion equation that simultaneously models the effects of demography and selection on pairs of derived alleles (Fig. 1). Using our method, we inferred, for the first time, the correlation of mutation fitness effects at the same site within proteins from triallelic nonsynonymous SNP data (Fig. 3). We found that the correlation coefficient is intermediate between completely uncorrelated and completely correlated. Early mutation-selection models of protein evolution made the unrealistic assumption that the fitness effects of multiple mutations occurring at the same site were identical [9]. More recent methods estimate selection coefficients for every possible amino acid at every site [10], but these complex models require a great deal of data [47]. Our model of correlated fitness effects is a useful intermediate complexity model.

We found strong quantitative agreement between the fitness effects correlation coefficient inferred from our population genomic inference and direct biochemical experiments (Fig. 4). Moreover, this agreement held across a wide range of model organisms, for genes that vary dramatically in function, and using several measures of fitness, suggesting that this correlation of mutation fitness effects is a fundamental property of protein biology, not species- or protein-specific. We also refined our analysis to biologically-relevant subsets of the data (Table 1). As expected, nonsynonymous pairs of similar derived amino acids show significantly higher correlation of fitness effects that dissimilar pairs. Although solvent accessibility and structural disorder did affect the marginal DFE (Table S3), we did not find a difference in fitness effects correlation between among these classes of sites (Table 1). Together, our results suggest that the fitness effects correlation we inferred is a nearly universal property of protein evolution, with important implications for modeling protein evolution.

## Methods

### Numerical solution to the triallelic PDE

The adjusted scaled selection coefficients 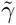 in Eq. 1 arise from competition between the two segregating derived alleles, dependent on their allele frequencies. If both derived alleles are at high frequency, they primarily compete against each other. For example, if their selection coefficients are roughly equal, even when that selection is strong, they will be effectively neutral when at high frequency. In general,

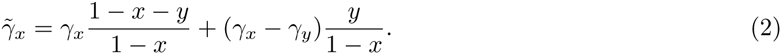

To integrate the diffusion equation forward in time, we used operator splitting to separately apply the non-mixed and mixed derivative terms each time step. We integrated the non-mixed terms using a conservative alternating direction implicit (ADI) finite difference scheme [48]. We integrated the mixed term using a standard explicit scheme for mixed derivatives. We used uniform grids in *x* and *y* with equal grid spacing Ä, so that grid points lie directly on the diagonal *x* + *y* = 1 boundary of the domain, which readily allowed the diagonal boundary to be absorbing. Although these integration schemes worked well in the interior of the domain, application at the diagonal boundary led to an excess of density being lost (Supplemental Text, Fig. S1). To avoid this excess loss of density near the diagonal boundary, we did not apply the ADI and mixed derivative schemes at the closest grid points to the diagonal boundary. Instead, at each time step we calculated the amount of density at each grid point that would fix along the diagonal boundary, and we directly removed that amount from the numerical density function and added it to the boundary.

To inject density into *φ* for new triallelic loci arising from mutation, at each time step we added density to the first interior rows of grid points based on the expected background biallelic frequency. For example, we added to the row of grid points *x* = Ä, *y* = Ä, 2Ä, … 1 − Ä with weight for point (Ä, *y*) proportional to the biallelic population allele density at frequency *y*.

To obtain the expected sample frequency spectrum *T* from the population frequency spectrum *φ*, we numerically integrated against the trinomial distribution with sample size *n*:

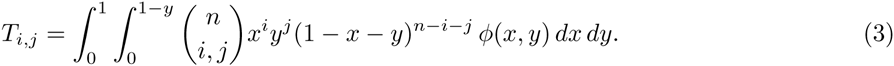

We estimated model parameters by maximum composite-likelihood. Likelihoods *L*(*D*|È) of the data *D* given the model parameters È were calculated by assuming that each entry in the observed triallelic frequency spectrum (*D_i,j_*) was an independent Poisson random variable with mean *T_i,j_* [49], where *T* is the expected triallelic frequency spectrum generated under È:

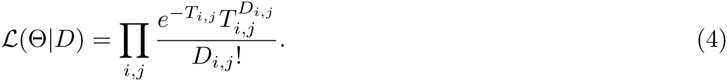

The overall population-scaled mutation rate was an implicit free parameter in our model fits. We calculate parameter uncertainties for each model fit using the Godambe Information Matrix [50]. Our code implementing these methods is integrated into ∂a∂i, available at http://bitbucket.org/RyanGutenkunst/dadi.

### Calculating frequency spectra under the joint DFE

To integrate over the bivariate DFE we used a logarithmically spaced grid with 50 grid points ranging from 10^−4^ to 2000 for negative *γ*, along with *γ* = 0 and *ã*+ = 39:9. We cached spectra for each possible pair (*ã_x_*, *ã_y_*), yielding 52^2^ cached spectra. A pair of selection coefficients (*ã_x_*, *ã_y_*) could fall into four quadrants depending on the sign of *ã_x_* and *ã_y_*. The overall frequency spectrum was calculated by summing over the weighted frequency spectra for each quadrant based on the DFE parameters *p*_+_ and *ñ*. The weights were 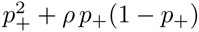 for both *ã_x_*, *ã_y_* > 0, (1 − *ρ*)*p*_+_(1 − *p*_+_) for one selection coe_cient positive and the other negative, and (1 − *p*_+_)^2^ + *ρ*(1 − *p*_+_)*p*_+_ for both *ã_x_*, *ã_y_* < 0. To integrate over the continuous distributions with either selection coefficient negative, we used the trapezoid rule. We approximated *ã* ∈ (0; 10^−4^) as effectively neutral and *γ* < −2000 as effectively lethal (Fig. S5).

### Genomic data

We extracted SNPs from the Drosophila Genome Nexus Data [19] and used Annovar [51] to determine the transcript and codon position of the coding SNPs. Ancestral states for each codon were determined using the aligned sequences of *D. melanogaster* (April 2006, dm3) and *D. simulans* (droSim1) downloaded from the UCSC genome database. We excluded loci with no aligned *D. simulans* sequence. We downloaded the reference transcript sequences from Ensembl Biomart [52] and used the ancestral states determined by the droSim1 alignment to determine the ancestral codon state.

### Mutational scanning data

We considered data from three mutational scanning studies [6, 31, 32]. Each assayed a different protein from a different organism using a different proxy for fitness. In all three experiments, the distribution of fitnesses was bimodal, with peaks of moderately and strongly deleterious mutations, although the relative sizes of these peaks differed markedly between experiments (Fig. S4A-C). To calculate the fitness correlation coefficient, we sampled a pair of mutually nonsynonymous mutations from each site in the protein (excluding mutations without reported fitness) and calculated the Pearson correlation of those fitnesses. The confidence intervals in Table 1 are 2.5% and 97.5% quantiles from 10,000 repetitions of this sampling. To visualize the correlations, we calculated the proportion of mutually nonsynonymous mutation pairs within each possible bin of joint fitness effects (Fig. S4D-I). Because our population-genetic analysis is not sensitive to strongly deleterious mutations, we focused our analysis on moderately deleterious mutations (shaded regions in Fig. S4A-C, joint distributions in Fig. S4D-F). For details on each data set, see Supplemental Text.

## Acknowledgments

This work was supported by the National Science Foundation (DEB-1146074 to RNG).

